# Novel chromosome organization pattern in actinomycetales–overlapping replication cycles combined with diploidy

**DOI:** 10.1101/094169

**Authors:** Kati Böhm, Fabian Meyer, Agata Rhomberg, Jörn Kalinowski, Catriona Donovan, Marc Bramkamp

**Author notes:** Corresponding author: Marc Bramkamp.

## Abstract

Bacteria regulate chromosome replication and segregation tightly with cell division to ensure faithful segregation of DNA to daughter generations. The underlying mechanisms have been addressed in several model species. It became apparent that bacteria have evolved quite different strategies to regulate DNA segregation and chromosomal organization. We have investigated here how the actinobacterium *Corynebacterium glutamicum* organizes chromosome segregation and DNA replication. Unexpectedly, we find that *C. glutamicum* cells are at least diploid under all conditions tested and that these organisms have overlapping C-periods during replication with both origins initiating replication simultaneously. Based on experimentally obtained data we propose growth rate dependent cell cycle models for *C. glutamicum*.

## Introduction

Bacterial chromosome organization is highly regulated, where replication coincides with segregation of sister nucleoids and is tightly coordinated with cell division (1). Cell cycle control mechanisms exist, which ensure constant DNA content throughout cell generations. In particular, the action of the key replication initiator protein DnaA is timed by various regulatory systems, for instance via the CtrA protein cascade in *Caulobacter crescentus* or SeqA in *Escherichia coli* (2-6). Upon replication initiation DnaA binds to the origin of replication *(oriC)* and mediates duplex unwinding prior to loading of the replication machinery (7,8). The two evolving replication forks migrate along the left and right arm of the circular chromosome towards the terminus of replication (*terC*), where FtsK-dependent XerCD recombinases resolve decatenated chromosomes as a final step (9,10). Replication usually takes place within defined cellular regions via stably assembled protein complexes, namely replisomes, of rather static or dynamic nature (11,12).

The bacterial cell cycle can be divided in different stages illustrated in Figure 1. The time of DNA-replication is termed C-period, which is followed by a time interval necessary for cell division executed by the divisiome (D-period). Several bacteria like *Mycobacterium smegmatis* and *C. crescentus* replicate their genome once within a generation, where C-periods are temporally separated from each other (13,14). At slow growing conditions a non-replicative state termed B-period precedes the C-period (not shown), thus the bacterial cell cycle resembles in some aspects the eukaryotic cell cycle (G1, S, G2 phases). Contrary to this, fast growing organisms such as *Bacillus subtilis, E. coli* and *Vibrio cholerae* can overlap C-periods during fast growth, a phenomenon termed multifork replication (15-17). Under these conditions a new round of replication is reinitiated before termination of the previous one. Therefore, generation times are considerably shorter than the duration of the C-period. However, only one round of replication is initiated per cell cycle and usually one C-period is completed at the time point of cell division (18). Many bacteria contain only one copy of the chromosome. However, several bacteria and archaea can have increased DNA contents due to oligo-or polyploidy (19). Polyploid cells harbor multiple, fully replicated chromosome copies throughout their life cycle, which has been frequently found in prokaryotes including certain gram-positive bacteria, proteobacteria, Deinococcales, cyanobacteria and also archaea (20-27).

**Figure 1.**
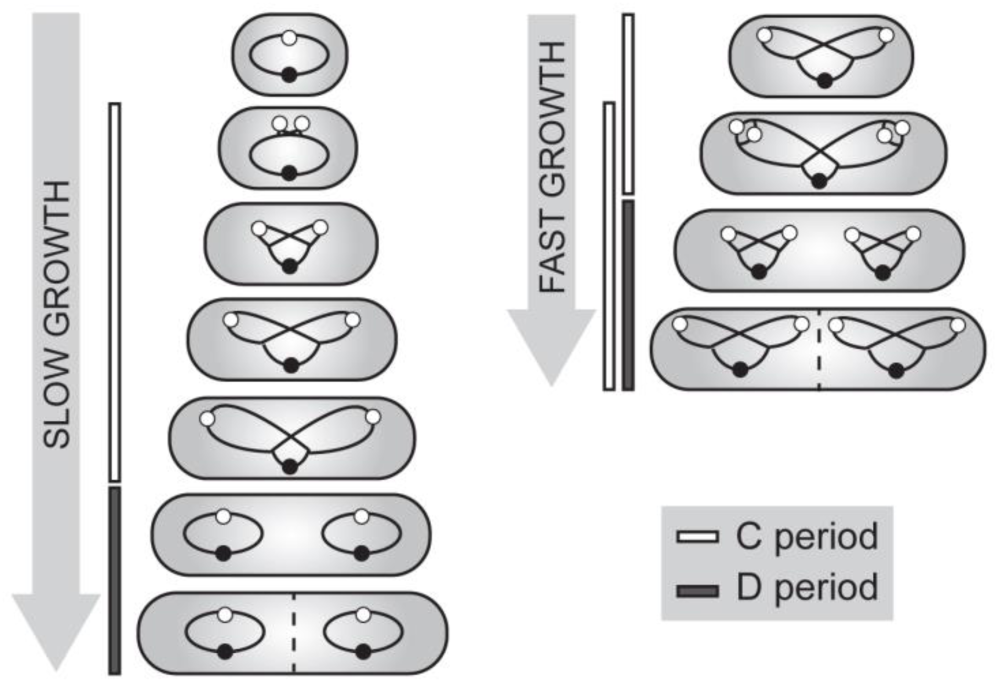
Schematic representation of bacterial replication cycles under slow (left) and fast (right) growth conditions. During slow growth conditions DNA replication (termed C-period) takes place within a single generation, followed by the interval between replication termination and completion of cell division (D-period). Fast growing bacteria with generation times of less than the C period, like *Bacillus subtilis* and *Escherichia coli*, undergo multifork replication, i.e. new rounds of replication are initiated before previous ones terminate. Chromosomes are indicated by black lines with *oriCs* and *terCs* as white and black circles.

Besides the distinct cell cycle modes chromosome localization patterns differ between model organisms. In a non-replicating, slow-growing *E. coli* cell the single chromosome is placed symmetrically with *oriC* and *terC* regions located at midcell and the replichores spatially separated to the two cell halves (28). Upon replication initiation the two sister chromosomes segregate bidirectionally to opposite cell halves with replisomes positioned at midcell (29,30). Finally, *oriC* and *terC* are confined to cell quarter regions. Contrary to this, the model organisms *C. crescentus, Vibrio cholerae* and *Pseudomonas aeruginosa* localize their nucleoids about the longitudinal axis with chromosome arms adjacent to each other (31-34). Sister replichores move to the opposite cell half with the segregated *oriC* facing towards the pole, mirroring the second chromosome at the transverse axis. The *oriC* region of *C. crescentus* and *V. cholera* is positioned by polar landmark proteins (35,36), where replisomes assemble and simultaneously move towards midcell in the course of replication (12,17). For the most part, *P. aeruginosa* places its replication machinery centrally (34). Finally, *B. subtilis* switches from the longitudinal chromosome organization to the *E. coli* “left-*oriC*-right” configuration during replication initiation (37).

The mitotic-like ParABS system has been identified as a driving force behind coordinated nucleoid partitioning for more than two third of bacterial species analyzed, with exceptions specifically within the phylum of γ-proteobacteria such as *E. coli* (38). This segregation mechanism involves components similar to the plasmid encoded *par* genes responsible for active segregation of low-copy-number plasmids (39). Thereby, the ParB protein binds a variable number of centromere-like DNA sequences called *parS* sites in *oriC*-proximity (40) and spreads along the DNA forming large protein-DNA complexes (41-43). Interaction of ParB with the Walker-type ATPase ParA mediates ATP-hydrolysis and thereby ParA detachment from DNA (44), driving apart the sister chromosomes as the protein interaction translocates the *oriC* towards the opposite cell half (33,45,46). The precise mechanism of the ParABS-mediated DNA segregation has been under debate, however, to date dynamic diffusion-ratchet and DNA-relay models are favored, where nucleoid and plasmid movement is mediated along a ParA gradient caused by local ParB-stimulated depletion of DNA-bound ParA (47-49). Deletion of this partitioning system has mild effects in *B. subtilis* and *V. cholera*e cells, but causes severe chromosome segregation defects in other organisms and is essential for viability in *C. crescentus* and *Myxococcus xanthus* (46,50-56).

Here we present the cell cycle and spatiotemporal organization of *oriCs* and replisomes in *C. glutamicum*, a rod-shaped polar growing actinobacterium. It is closely related to pathogens like *C. diphteriae* and *M. tuberculosis*, the latter being amongst the top ten causes of fatal infections worldwide (57). Besides this, *C. glutamicum* is of great economic importance as an amino acid and vitamin producer and extensive efforts in metabolic engineering are being carried out concerning metabolite production and yield increase (58). Although, its metabolism is one of the best studied amongst model organisms the underlying cell cycle parameters and chromosome organization patterns had so far not been analyzed in detail. *C. glutamicum* relies on a ParABS system to segregate their nucleoids prior to cell division (51,59,60). Chromosome segregation influences division site selection and, hence, growth and chromosome organization are tightly coupled in *C. glutamicum* (59). This may in part explain why protein machineries that have been described in various bacterial species like the Min system or a nucleoid occlusion system, both being involved in division septum placement are absent in *C. glutamicum* (61).

In this study, we tracked *in vivo* fluorescently labeled centromer-binding protein ParB and replisome sliding clamp DnaN to investigate spatiotemporal *oriC* and replisome localization throughout the cell cycle. Fluorescence microscopy and single cell tracking via time lapse analysis revealed remarkably high *oriC* and replisome numbers during fast growth, suggesting multiple chromosomes and several simultaneous replication events per cell. Initially cells possess two polar *oriC*-ParB cluster, whereby upon replication sister *oriC*s segregate towards midcell positions. Additionally, the length of replication periods as well as the overall DNA content was determined for different growth conditions by marker-frequency analysis and flow cytometry, thereby allowing the formulation of complete cell cycle models. Our data suggest diploidy and overlapping C-periods in *C. glutamicum* and therefore, give new insights in replication coordination within the group of actinobacteria.

## Materials and Methods

### Oligonucleotides, plasmids and bacterial strains

All primers, plasmids and bacterial strains used in this study are listed in Table S1 and S2, respectively.

### Strain construction

Integration plasmids were constructed with 500 bp homologous regions upstream and downstream of the 3’ end of the gene to be labeled with a fluorophore sequence in between. For pK19mobsacB-parB-eYFP and pK19mobsacB-parB-mCherry2 plasmids the upstream and downstream fragments were PCR amplified from the *C. glutamicum* genome using the primer pairs ParB-Hind-up-F/ ParB-Sal-up-R and ParB-XbaI-D-F/ ParB-Bam-D-R. eYFP or mCherry2 sequences were amplified using primer pairs eYFP-SalI-F/ eYFP-XbaI-R or mCherry2-SalI-F/ mCherry2-XbaI-R. In order to construct pK19mobsacB-DnaN-mCherry upstream and downstream homologous regions were amplified via primer pairs DnaN-Hind-up-F/ DnaN-SphI-up-R and DnaN-XbaI-D-F/ DnaN-BamHI-D-R. The mCherry sequence was amplified using mCherry-SalI-F/ mCherry-XbaI-R primer pairs. The resulting PCR fragments were digested with the respective restriction enzymes and consecutively ligated into pK19mobsacB vectors. Plasmid cloning was performed using DH5α. *C. glutamicum* were transformed via electroporation and selected for integration of the fluorophore as described before (62). To confirm the allelic replacements *parB::parB-eYFP* and *parB::parB-mCherry2* colony-PCR was carried out using primers ParB-N-ter-SalI-F and ParB-PstI-800D-R and for the allelic replacement *dnaN::dnaN-mCherry* primers DnaN-N-ter-F and DnaN-Bam-700D-R were used.

### Growth conditions and media

*B. subtilis* and *E. coli* cells were grown in Luria-Bertani (LB) medium supplemented with Kanamycin when appropriate at 37°C. *C. glutamicum* cells were grown in brain heart infusion (BHI, Oxoid™) medium, BHI medium supplemented with 4% glucose or in MMI medium (63) supplemented with 4% glucose, 120 mM acetate or 100 mM propionate as indicated in the text at 30°C. For growth experiments in BHI supplemented with glucose or MMI media cells were inoculated in BHI and diluted in growth media overnight for pre-cultivation. The next morning cultures were diluted to OD_600_ 1. Growth in BHI preceded an overnight inoculation step; cultures were re-diluted to OD_600_ 0.5 the next morning. For replication runouts exponentially growing C. *glutamicum* or *B. subtilis* cells were treated with 25 μg/ml or 200 μg/ml chloramphenicol for 4+ h.

### Fluorescence microscopy and life cell imaging

Fluorescence microscopy was carried out using an Axio-Imager M1 fluorescence microscope (Carl Zeiss) with an EC Plan Neofluar 100x/ 1.3 oil Ph3 objective and a 2.5 x optovar for automated image analysis. Filter sets 46 HE YFP (EX BP 500/25, BS FT 515, EM BP 535/30) and 43 HE Cy 3 shift free (EX BP 550/25, BS FT 570, EM BP 605/70) were used for fluorescence detection of eYFP and mCherry or mCherry2 protein fusions. DNA was stained using 1 μg/ml Hoechst 33342 (Thermo Scientific). For life cell imaging cells at exponential growth were re-diluted in BHI medium to OD_600_ 0.01 and loaded in a microfluidic chamber (B04A CellASIC®, Onix) at 8psi for 10 sec; for nutrient supply 0.75 psi were applied. Time lapse microscopy was performed using a Delta Vision Elite microscope (GE Healthcare, Applied Precision) with a standard four color InsightSSI module and an environmental chamber heated to 30°C. Images were taken with a 100×/ 1.4 oil PSF U-Plan S-Apo objective and mCherry (EX BP 575/25, EM BP 625/45) or YFP (EX BP 513/17, EM BP 548/22) specific filter sets were used for fluorescence detection (50% transmission, 0.3 sec exposure). Images were taken in 5 min intervals. For data analysis FIJI software (64) was applied; cell length measurements were acquired manually.

### Marker frequency analysis

Genomic DNA was isolated from *C. glutamicum* or *B. subtilis* cells in exponential or stationary growth phases. DNA proximal to the origin or terminus (see text) was amplified by qPCR using a qPCR Mastermix (KAPA SYBR®FAST, Peqlab) according to manufacturer’s protocol. Each experiment was performed in technical triplicates. Oligonucleotides used are listed in Table S1 and results were analyzed using the 2^-ΔCT^ method (65).

### Flow cytometry analysis

Culture samples were fixed 1:9 (v/v) in 70% ethanol and stored at 4°C until use. Cells were pelleted at 5000 rpm for 5 min and washed once in phosphate-buffered saline (PBS). The DNA staining procedure was adopted from a protocol described before (66). Samples were preheated to 37°C and stained in SYBR^®^ Green I (Invitrogen) with a final dilution of 1:10000 for 15 min and consequently diluted in PBS. Flow cytometry analysis was performed using a BD Accuri C6 (BD Biosciences) with a 488 nm blue laser. The measurements were conducted at a flow rate of 10 μl/min with an acquisition threshold set to 700 on FL1-H and a rate of events per second less than 5000. At least 200000 events per sample were collected. Data were analyzed by plotting samples as histograms vs the green channel (FL1-A, EM BP 533/30) at log scale. All experiments were performed with biological triplicates.

In order to calibrate the DNA measurements of different growth conditions *B. subtilis* cells were used as an internal standard. A replication runout of *B. subtilis* cells grown in LB medium gave rise to cells with mainly 4 or 8 fully replicated chromosomes (67). Prior to ethanol fixation the cell wall was stained via strain-promoted alkyne-azide cycloaddition (SPAAC). In short 5 mM 3-azido-D-alanine (Baseclick GmbH), which incorporates into the cell wall, was added to the culture during the time of replication runout. Cells were washed in PBS, incubated with 10 μm DBCO-PEG_4_-5/6-Carboxyrhodamine 110 (Jena Bioscience) at 30 °C for 20 min in the dark and subsequently washed three times in PBS + 0.1% Tween 80. This standard was included with *C. glutamicum* cells during incubation with 1 μg/ml Hoechst 33342 DNA stain. Flow cytometry was performed with a FACSAria II (Becton Dickinson) using a 488 nm blue laser and a 355 nm UV laser and appropriate filter sets. 50000 events were collected per sample. Blots of DNA content vs. the green channel were used to identify *B. subtilis* subpopulations and *C. glutamicum* chromosome numbers were assessed in accordance with the standard in histograms vs. DNA amount.For data analysis BD Accuri C6 software (BD Biosciences) or FlowJo software (Tree star, Inc.) were applied.

### Analysis of the cell cycle

C-and D-periods were determined via equations relating to DNA amount per cell in exponential cultures (67,68), which were adapted to the *C. glutamicum* cell cycle model with double the number of chromosome equivalents at any time. Since only every second initiation is followed by a cell division the average *oriCs* per cell (Ī) are defined as shown below. The term for the average *terCs* per cell (T̅) was adjusted accordingly, where т is the doubling time:

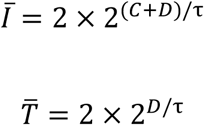

Hence, the D period was calculated as shown below; the average number of *oriCs* (I) per cell (N) was resolved by flow cytometry:

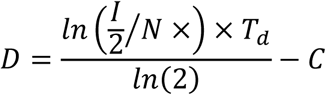

The equation for determination of C periods does not change upon assumptions made above, where the *oriC* to *terC* ratio (I/T) was determined by marker frequency analysis:

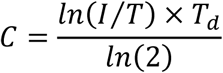

### Statistical analysis

ANOVA and post hoc tests were performed using R (69); correlation coefficients, ANCOVA and linear regressions were calculated using Excel and Graph Pad Prism (GraphPad Software).

## Results

### Origin numbers correlate with cell length in a ParA-independent way

We have shown before that the *C. glutamicum* partitioning protein ParB localizes at the origin regions of the chromosome close to the cell poles (51). However, in depth studies of spatiotemporal chromosome organization were still missing. Therefore, we aimed to reanalyze *oriC*-ParB complexes microscopically by time resolved life cell imaging. In order to label origin regions the native chromosomal *parB* locus of *C. glutamicum* RES167 wild type (WT) or in a *ΔparA* mutant was replaced by *parB-eYFP*, resulting in strains with typical cell morphology and growth phenotypes (Fig. 2A, Fig. S1). No cleavage products of ParB-eYFP were detectable (Fig. S2), suggesting that fluorescent signals faithfully reflect ParB localization. Microscopic analysis revealed a correlation of ParB-eYFP foci numbers with cell length in both strains (Fig. 2B). In the WT background between one and five foci were detected. The *parA* deletion mutant has variable cell lengths and anucleate minicells (not taken into account) were observed. Up to twelve ParB foci were present in cells of the *parA* deletion mutant. Notably, most of the ParB foci in the *parA* knockout strain were less fluorescent compared to the situation in wild type cells, suggesting that lack of ParA causes problems in the ParB assembly at the origin. Chromosome segregation defects upon *parA* deletion do not markedly affect the high correlation of *oriC*-ParB foci number to cell length. However, since linear regression models yield significant differences between WT and the *parA* deletion strain (Fig. 2B) a loss of ParA might cause slight over-initiation of chromosome replication.

**Figure 2.**
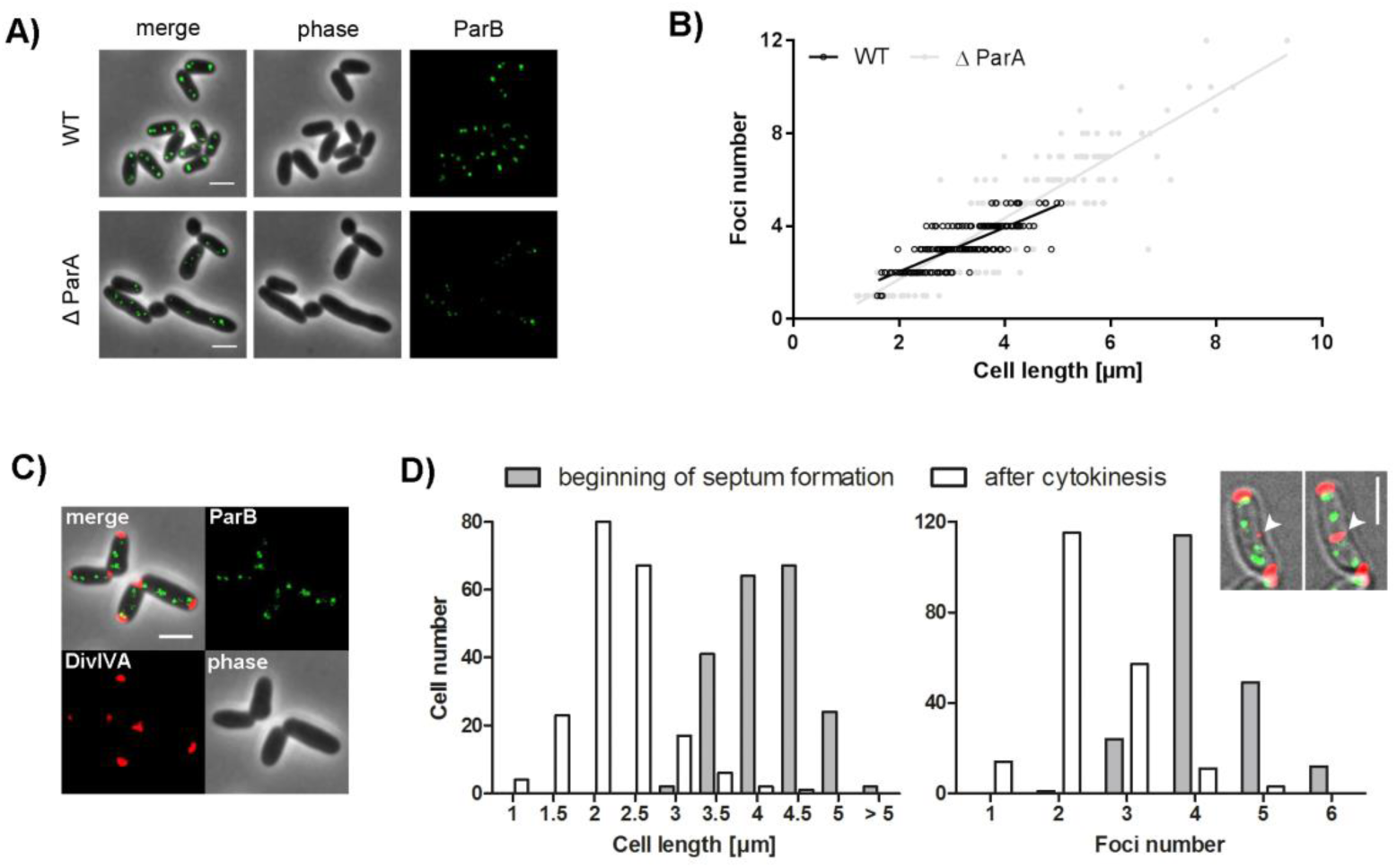
Determination of the *C. glutamicum oriC* number and correlation to cell length. **A)** Subcellular localization of ParB-eYFP in representative *parB⸬parB-eYFP* wild type (WT) and Δ*parA* cells. Shown are overlays between phase contrast images and eYFP fluorescence (merge) and separate channels (phase, ParB). Scale bar, 2 μm. **B)** ParB foci number depend on cell length in a ParA independent way. In wild type strain (WT) one to five foci and in Δ*parA* strain one to twelve foci were observed (n=400). Linear regression lines are shown r(wild type)=0.80, r(Δ*parA*)=0.88; slopes are not equal (ANCOVA, F(1, 396)=16.10, p <.0001). **C)** Still image of *C. glutamicum parB⸬parB-eYFP divIVA⸬divIVA-mCherry* cells. Depicted are phase contrast (lower right), DivIVA (lower left) and ParB-eYFP fluorescence (upper right) and the overlay of all channels (upper left). Scale bar, 2 μm. **D)** Time-lapse microscopy of DivIVA-mCherry and ParB-eYFP coexpressing strain reveals distribution of cell length and ParB-eYFP cluster number at the beginning of septum formation and after cell division (n=200). The microscopy images of a single cell exemplify the time lapse of septum formation (white arrowhead) tracked by DivIVA-mCherry reporter. Scale bar, 2 μm.

In order to determine the maximal number of origins per cell precisely, an additional allelic replacement of *divIVA* by *divIVA-mCherry* was carried out (Fig. 2C). With this construct cell division can be monitored by DivIVA localization before the cells walls of the daughter cells are separated, because DivIVA efficiently accumulates at the septal membrane. This strain revealed WT-like growth rates and cell length distributions (Fig. S1), suggesting that DivIVA-mCherry and ParB-eYFP are functional. DivIVA localizes to the cell poles as well as newly formed septa and therefore, is a suitable marker for completed cell division (60). Analysis revealed an average of four ParB foci prior to completed septation at an average cell length of 3.94 μm, however, up to six foci per cell could be determined (Fig. 2D). Newborn cells with an average length of 2.01 μm mostly contained 2 foci. Origin numbers and cell size were relatively variable, as division septa are often not precisely placed at midcell in *Corynebacteria.* Using the DivIVA reporter to judge ParB foci number per daughter cell resulted in similar origin numbers per cell compared to using physical separation of daughter cells as a mark for cell division. Notably these analyses revealed unexpectedly high *oriC*-ParB cluster numbers that hint to the ability of *C. glutamicum* to either undergo multifork replication and/or to harbor multiple fully replicated chromosomes per cell.

### Spatiotemporal localization pattern of ParB-origin complexes

For analysis of chromosome arrangement during one cell cycle, life cell imaging was performed using strain *C. glutamicum parB::parB-eYFP* (Fig. 3A, Movie S2). Single cells with two ParB cluster were tracked over one generation time and foci positions were determined relative to the new pole, revealing distinct origin localization patterns throughout the cell cycle (Fig. 3B). New born cells contain two ParB foci stably located close to the cell poles. Newly replicated origins segregate from cell poles towards midcell. Often we observed the appearance of a new ParB cluster at either the new or old pole before a forth focus separates from the opposite ParB spot. No bias in timing of origin replication and segregation between old and new cell pole could be detected, as indicated by mean localizations of the third up to the fifth focus appearing around mid-cell positions. A timeline over one generation cycle shows the continuous increase of newly formed segregating origins (Fig. 3C). Already after completion of half a cell cycle (~30 min) around half of the cells contained four or five ParB foci; this ratio further increased during growth progression. In order to corroborate these findings we used automated analysis of still microscopy images to confirm the spatiotemporal *oriC*-ParB cluster localization (Fig 3D). Therefore, we programmed a Fiji software plug-in termed Morpholyzer that allows automated cell detection and analysis of fluorescent profiles (see material an methods). High ParB fluorescence intensities were detected close to the poles for all cells measured, suggesting stable origin anchoring at the cell poles. Segregation of sister *oriC*-ParB cluster towards midcell positions could be detected after around one fourth of the cell cycle (Fig. 3D). Similar dynamics of origin localization and segregation patterns have been characterized before for *M. xanthus*, *C. crescentus*, *M. smegmatis* and *V. cholera* (33,70-73). Upon ParA deletion the time dependent increase of origin numbers became less distinct due to large cell length variations directly after cytokinesis (Fig. 3E). As a consequence, already in large newborn cells multiple *oriC-*ParB complexes were present. Analysis of still images further revealed a disrupted ParB-origin pattern in *parA* deletion strains compared to the coordinated cellular origin movement in WT cells (Fig. 3F). Fluorescent foci were detected all along the longitudinal cell axis without clear sites of preference, underlining the crucial role of the partitioning protein ParA in polar and septal positioning of *oriC*-ParB cluster in *C. glutamicum*.

**Figure 3.**
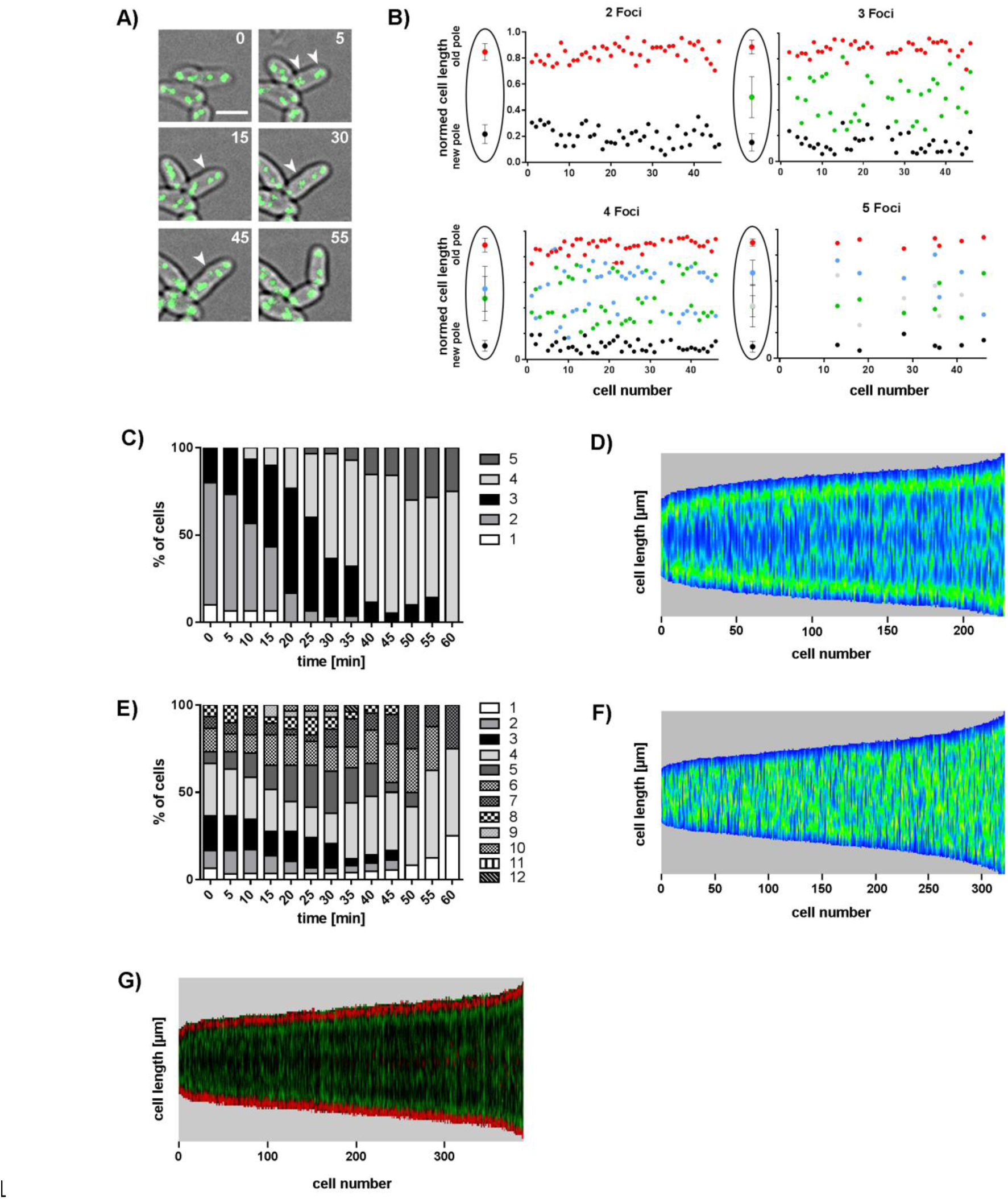
*OriC* localization pattern during cell cycle progression. **A)** Occurrence of newly formed ParB-eYFP clusters in the course of cell elongation. Still images show a time series of a typical *C. glutamicum* wild type cell with initially two ParB spots. Three further foci appear over time (white arrowheads); time points are indicated in min (top right corner). Scale bar, 2 μm. **B)** Time lapse single cell analyses reveal *oriC*-ParB complex positions along the long cell axis at each moment in time a new ParB-eYFP spot occurs. A third, fourth and eventually a fifth focus separate from the two initial ParB clusters located close to the cell poles and move towards midcell positions. Cells are aligned with the old pole facing upwards; cell lengths are normed to 1 (n=46). Schemes as shown to the left illustrate average ParB-eYFP foci positions ± standard deviation. **C)** Time dependent increase of ParB clusters per cell. Fraction of cells with 1-5 spots are depicted for each time point (n=30). **D)** ParB-eYFP pattern along the cell axis in dependence of cell length in wild type *C. glutamicum.* Automated image analysis of still microscopy images sorted by cell length with high fluorescence intensities displayed in green (n>200). **E)** Counts of ParB-eYFP spots over time in *C. glutamicum ΔparA.* Percentages of cells with 1-12 ParB foci were determined for each time point (n=30). **F)** Random ParB-eYFP distribution along the longitudinal cell axis in relation to its length in Δ*parA* mutant strain. Automated analysis of still images with high fluorescence intensities displayed in green (n>300). **G)** Timing of replication initiation is similar at *oriC* of old and young cell pole. Automated image analysis of *C. glutamicum parB⸬parB-eYFP divIVA⸬divIVA-mCherry* fluorescence pattern sorted by cell length with the old cell pole (high polar DivIVA-mCherry signal) facing downwards. ParB-eYFP (green) and DivIVA-mCherry (red) fluorescence are illustrated in one kymograph.

### Uniform timing of replication initiation at old and young cell poles

*C. glutamicum* cells grow asymmetric by unequal rates of peptidoglycan-synthesis (PG) at the cell poles. The old cell pole synthesizes more PG compared to the young pole (74) due to cell cycle dependent dynamics of DivIVA accumulation (own unpublished data). As *C. glutamicum* DivIVA interacts with the origin-attached ParB protein (60) an impact of the DivIVA level on chromosome replication or segregation timing was investigated. Using automated analysis of cells co-expressing ParB-eYFP and DivIVA-mCherry old cell poles were identified by higher DivIVA fluorescence intensities and aligned accordingly. The fluorescence profile visualized in Figure 3G reveals synchronous origin movements towards the newly formed septum from both old and new cell poles, as suggested before by time-lapse analysis (Fig. 3B). Therefore, timing of chromosome replication seems to be uncoupled from the assembly of the cell wall synthesis machinery despite ParB-DivIVA interaction.

### Replisome tracking reveals multiple replication forks and variable origin cohesion times

*In vivo* characterization of replisome dynamics was carried out using a reporter strain in which the native locus of helicase sliding clamp *dnaN* was replaced by a *dnaN-mCherry* fusion construct (Fig. 4A). Cell length distribution and growth of the DnaN-mCherry expressing strain resembles the WT situation (Fig. S1) and the presence of full-length DnaN-mCherry protein could be confirmed via western blot (S2), suggesting that the localization patterns are not due to free fluorophores or degraded protein. In order to track the pattern of replication timing automated analysis of fluorescence microscopy images was applied on cells grown in BHI (Fig. 4B). In growing cells high DnaN-mCherry signals were observed in a wide range around midcell. At the end of the first third of the generation time a fluent transition of replication termination around midcell towards formation of newly formed replication hubs in cell quarter positions took place. This large scale microscopy analysis clearly indicates that new rounds of replication initiation cannot be temporally separated from the previous ones. C-periods follow each other at short intervals or might even overlap during fast growth conditions. Notably, single cell analysis can show that replisomes are formed at polar or septal *oriC*-ParB complexes and gradually move away from the origins towards midcell (Fig. S3). Such a DnaN fluorescence pattern is not immediately obvious in kymographs, presumably due to variable timing of replication initiation between cells of similar size. The movement of replication forks observed by life cell imaging appeared to be highly dynamic (Movie S2). During life cell imaging a progressive increase of DnaN-foci over one generation time was observed (Fig. 4C). Initially two replication forks were counted for most of the cells; the number further increased to up to six foci per cell throughout the cell cycle indicating three or more simultaneous replication events per cell. In order to further analyze replication initiation and progression in dependence of origin localization a dual-reporter strain expressing ParB-eYFP and DnaN-mCherry was constructed. WT-like growth and cell lengths for this strain were confirmed (Fig. S1). The dependence of ParB-eYFP and DnaN-mCherry foci numbers on cell length is shown in Figure 4D. On average less DnaN than ParB spots were counted per cell, as indicated by regression lines. However, the replication fork number could have been underestimated due to frequent merging of forks initiated from the same origin. These results reveal simultaneous replication events of several chromosome equivalents per cell. Furthermore, a moderate correlation between number of *oriC-ParB* clusters and replication forks per cell could be determined. Using life cell imaging cohesion periods of sister origins were analyzed, which are defined as the time between the formation of a new DnaN-mCherry spot in colocalization with an *oriC-ParB* cluster and the subsequent splitting of the latter into two distinct fluorescent signals (Fig. 4E). Time frames between replication initiation and sister origin segregation are illustrated in Figure 4F, revealing cohesion periods measured between 5 up to 80 min with an average of 36 min. The mean interval length is comparable to the 40 min origin colocalization period in fast growing *E. coli* cells determined before (75). As origin colocalization periods appear to be highly variable, a tight regulation for their cohesion might be absent in this organism. Replication initiations in the mother generation could frequently be observed with origin splitting only in subsequent generations as exemplified in Figure 4F. Notably, new rounds of chromosome replication were initiated at polar as well as at midcell positioned origins (Fig. S3).

**Figure 4.**
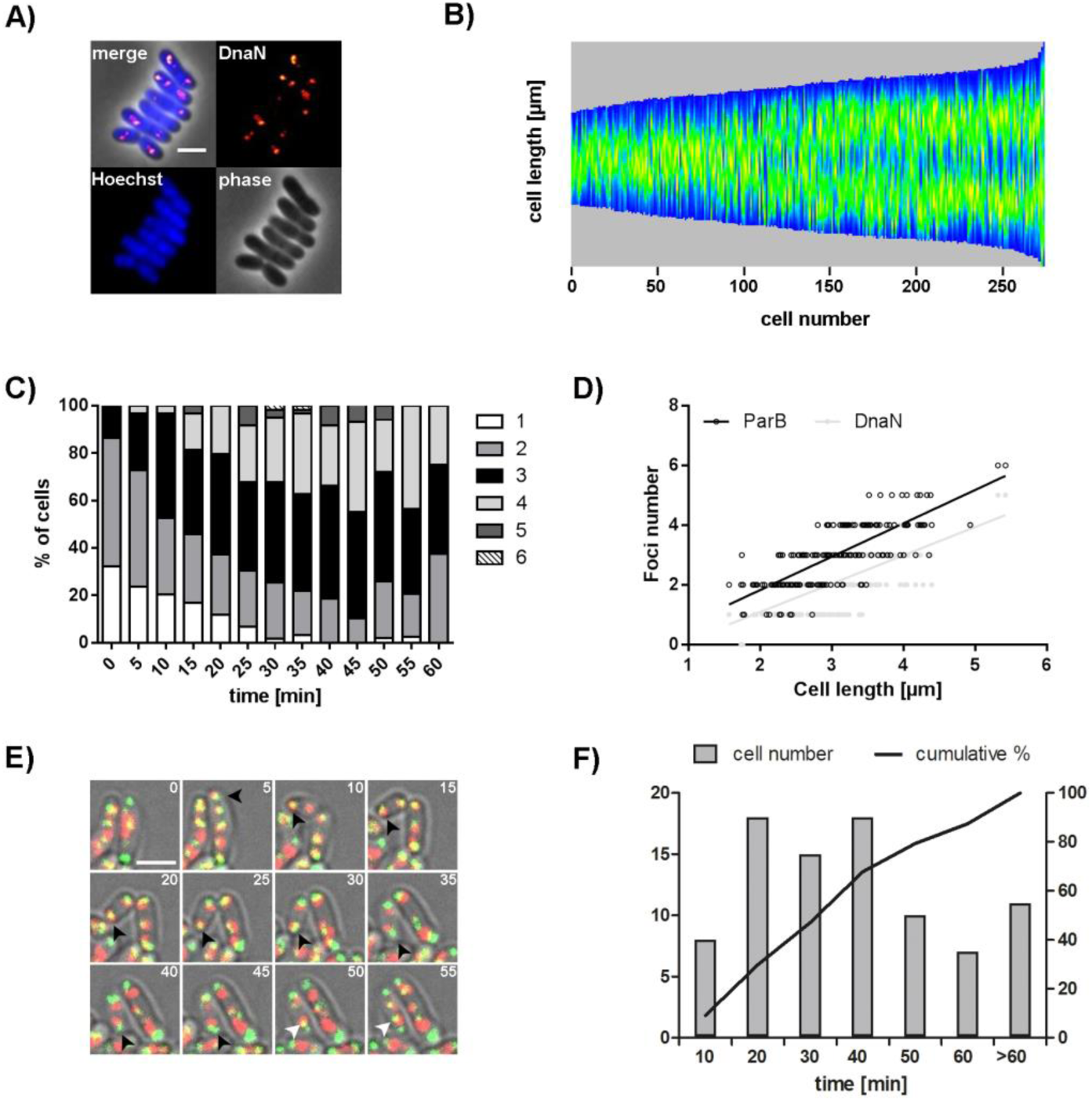
Dynamic localization of multiple replisomes in *C. glutamicum*. **A)** Localization of replisomes in *dnaN⸬dnaN-mCherry* cells. Shown is the overlay (merge) of DnaN fluorescence (red) and DNA stained with Hoechst (blue) with the phase contrast image and separate channels (DnaN, Hoechst, phase). Scale bar, 2 μm. **B)** Timing of replication along the cell axis. Automated analysis of still images with DnaN-mCherry fluorescence. Cells are sorted by cell length and DnaN-mCherry fluorescence is shown as heat map (blue to orange) (n>250). **C)** Replisome number per cell varies within one cell cycle. Fraction of cells with 1-6 DnaN spots were determined for each time point (n=59). **D)** ParB and DnaN foci number in relation to cell length in *C. glutamicum parB⸬parB-eYFP dnaN⸬dnaN-mCherry* (n=200). Linear regression lines are shown, r(ParB-eYFP/DnaN-mCherry)=0.65. **E)** Time frames of replication initiation until segregation of sister *oriCs.* Time series showing movement of ParB-eYFP and DnaN-mCherry foci (green and red, overlay in yellow) in *parB⸬parB-eYFP dnaN⸬dnaN-mCherry* cell. Images were taken in 5 min time-frames as indicated (top right corner). At time point 5 a replisome forms at polar *oriC* (black arrowheads); sister *oriCs* separate at time point 50 (white arrowheads). Scale bar, 2 μm. **F)** Variable cohesion periods of sister *oriCs.* Shown is the distribution of *oriC* colocalization times analyzed by time lapse microscopy together with the cumulative skew of sample data (n=88).

### Overlapping replication periods allow for fast growth

The observation of multiple DnaN and ParB foci suggest the possibility that *C. glutamicum* is able to initiate new replication rounds before finishing the ongoing replication. To test this hypothesis, we applied marker frequency analysis to investigate growth rate dependent replication patterns of *C. glutamicum. OriC/ter* ratios were determined for cells grown in three different media (BHI, BHI+Gluc or MMI) allowing for fast, intermediate or slow growth (Fig. S4A). Data from qPCR experiments using markers of origin and terminus proximal regions prove a growth rate dependency of *oriC/ter* ratios (Fig. 5A). As control we analyzed *B. subtilis* cells an organism with clear multi-fork replication. Exponentially grown *B. subtilis* cells result in *oriC/ter* ratios considerably above of 2 (Fig. S4C). Analysis of exponentially growing *C. glutamicum* cells cultured in BHI and BHI medium supplemented with glucose yielded mean *oriC/ter* ratios of 2.4 and 2.2, indicating an overlap of replication periods. Under slow growth conditions in MMI medium the mean *oriC/ter* ratio of 1.7 did not significantly differ from values obtained from cells in stationary growth phases. Upon antibiotic treatment leading to replication run-outs (by inhibiting replication initiation yielding fully replicated chromosomes) the ratios dropped to values close to one (data not shown). These results were further supported by whole genome sequencing of cells grown in BHI and MMI medium (Fig. 5B). Sequencing coverages revealed a symmetric progression of replication forks on both arms of the chromosome for all conditions tested. During exponential growth in BHI the mean coverage of origin regions was around 2.1 fold the one measured for terminus proximal regions. The *oriC/ter* ratio dropped to 1.5 for log-phase cells grown in MMI medium as well as for cells of stationary growth phases. These results hint to a fraction of cells, which did not complete replication at stationary growth. Likewise, active replication forks were observed in around 24 % of stationary *M. smegmatis* cells (73). Marker frequency analysis and genome sequencing results point to growth rate dependent replication cycles in *C. glutamicum* cells, where the timing of a new initiation precedes termination of the previous replication event in order to enable fast growth.

**Figure 5.**
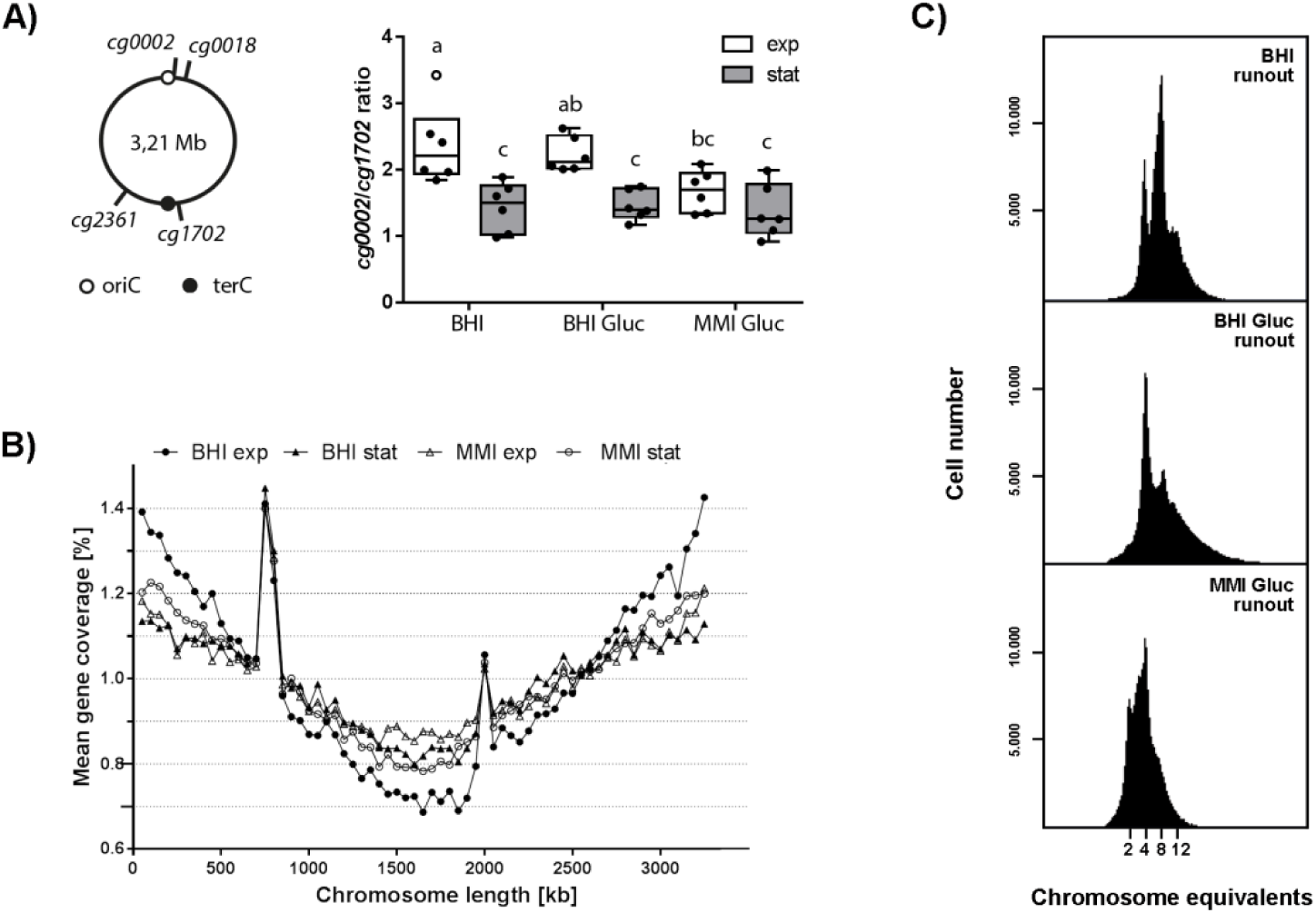
Timing of DNA replication initiation and determination of *oriC* numbers per cell. **A)** Marker frequency analysis of *oriC* and terminus regions. Right: Schematic representation of chromosomal positions of *oriC* and *terC* proximal marker genes applied. Left: *oriC* to terminus ratios of the wild type strain grown under different growth conditions were determined by frequency analysis of markers *cg0002* and *cg1702* (see S4B for *cg0018/cg2361* ratios). Right: Cells were grown in BHI, BHI supplemented with glucose or MMI supplemented with glucose. Samples were taken in the exponential (white boxes) and stationary growth phases (gray boxes). Results are shown as boxplots with medians indicated as solid lines and whiskers of 1.5 × interquartile range (n=6). An ANOVA yielded significant variation among conditions growth phase (F(1, 30)=28.00, p<.0001) and medium (F(2, 30)=3.43, p <.05). Letters indicate significant differences between data sets determined by Post-hoc Bonferroni analysis at p<.05. **B)** Whole genome sequencing. Genomic DNA of *C. glutamicum* wild type grown in BHI or MMI supplemented with glucose was isolated in exponential and stationary growth phases. Data were analyzed by Illumina MiSeq^^®^^ shotgun sequencing and mapped to the *C. glutamicum* ATCC 13032 genome sequence (GeneBankID: BX927147.1). Data are displayed as the mean gene coverage in % of each 50-kb sliding window of the total mean coverage per sample. Note that the RES167 strain used in this study features besides a lack of the phage island (cg1981-cg2034) an *ISCg14* mediated 5 × tandem amplification of the *tus* locus (peaks at approx. 750 and 2000 kb positions); both loci were excluded from data analysis. Stable replication progression is evidenced by frequency of genes between *oriC* (located at 0 kb) and terminus regions (at approx. 1.6 Mb). **C)** Chromosome numbers per cell determined by flow cytometry after replication runout in BHI, BHI supplemented with glucose or MMI supplemented with glucose. Depending on growth conditions between 2 and 12 chromosomes were detected.

### *C. glutamicum* cells contain multiple chromosome equivalents at varying growth rates

As shown by marker frequency analysis *oriC/ter* ratios are considerably higher in fast growing compared to slow growing cells. In order to more precisely verify the DNA content per cell flow cytometry was applied. To this end, replication run-out of WT cells cultured at varying growth rates were performed and nucleoids were fluorescently stained with SYBR^®^ Green I dye. DNA histograms show the number of fully replicated chromosomes, which equal the cellular origin numbers at the time point of antibiotic treatment (Fig. 5C). Absolute DNA content was assigned according to an internal standard (Fig. S5). Fast and intermediate growth conditions gave rise to mainly four and eight chromosomes per cell and a smaller fraction of cells contained ten and twelve chromosomes. Slow growth conditions yielded mainly cells containing either two or four and a small subpopulation containing eight chromosomes. Our data result in an average chromosome number per cell of 5.90, 5.17 and 3.85 from highest to lowest growth rate, respectively. Strikingly, the number of chromosome equivalents determined by flow cytometry is considerably higher than expected from marker frequency results and monoploid cell fractions were absent for all growth conditions tested in exponential phases. Flow cytometry results are further supported by fluorescence microscopy analysis of WT *parB::parB-eYFP* cells cultivated at several different growth rates. Cells with less than two ParB-foci per cell were almost absent under all condition tested. In addition, ParB-origin cluster numbers could possibly be underestimated due to potential origin cohesion (Fig. S6). Thus, we conclude that *C. glutamicum* is at least diploid with two chromosomes attached via the centromeric *oriC*-ParB nucleoprotein complex to the cell poles. Overinitiation of DNA replication leads to multi-ploid cells under fast growth conditions.

### Growth rate dependent cell cycle models

Cell cycle parameters derived from marker frequency and flow cytometry analysis (Table 1) allowed us to formulate complete cell cycle models at different growth conditions for *C. glutamicum*. A C-period of 78 min was determined for cells grown in BHI medium, whereas intermediate and low growth rates are associated with slightly longer replication periods of 96 and 97 min. These values result in a DNA replication speed of around 340 bases/ sec, which is in the range of replication speeds reported for *C. crescentus, M. xanthus* or *M. smegmatis* (Table 2 and references therein). D-period equations reported before (67) yielded time intervals longer than the doubling time for each of the three growth conditions analyzed indicating the presence of two fully replicated chromosomes per newborn cell. Since we define the D-period as time interval from replication termination that took place in the current generation until subsequent cell division D period calculation was adapted to a diploid organism (see Material and Methods). Those timeframes remained relatively unaltered with 20, 18 and 26 min from fast to slow growth rates, respectively.

**Table 1.** Overview of *C. glutamicum* cell cycle parameters at distinct growth rates.

| Growth medium | $\mu$<br>[1/h] | $T_d$ [min] | *oriC/terC* | *oriC/cell* | C [min] | D [min] |
| --- | --- | --- | --- | --- | --- | --- |
| BHI | 0.66 | 63 | $2.36 \pm 0,54$ | $5.90 \pm 0,03$ | 78 | 20 |
| BHI Gluc | 0.50 | 83 | $2.23 \pm 0,21$ | $5.17 \pm 0,02$ | 96 | 18 |
| MMI Gluc | 0.32 | 130 | $1.68 \pm 0,28$ | $3.85 \pm 0,16$ | 97 | 26 |

**Table 2.** Speed of DNA replication forks summarized for different model organisms.

| Organism | Genome size<br>[Mb] | C- period<br>[min] | Replication speed<br>[bases/sec] | Reference |
| --- | --- | --- | --- | --- |
| *Corynebacterium glutamicum* | 3.21 | 78 | 340 | this work |
| *Mycobacterium tuberculosis* | 4.42 | 660 | 50 | (85) |
| *Mycobacterium smegmatis* | 6.99 | 140 | 400 | (72,73) |
| *Myxococcus xanthus* | 9.14 | 200 | 380 | (70,86) |
| *Caulobacter crescentus* | 4.02 | 95 | 350 | (87) |
| ***Vibrio cholerae*\*** | 2.96 | 30-50 | 490-820 | (17,88) |
| ***Bacillus subtilis*** | 4.22 | 50-60 | 600-700 | (67,89) |
| ***Escherichia coli*** | 4.64 | 40-200 | 200-1000 | (15,67,90-92) |

\*Shown are parameters for chromosome I only.

Growth rate dependent cell cycle modes are illustrated for fast and slow growth conditions (Fig. 6). Newborn cells localize their origins in two clusters at polar positions with chromosomes being arranged longitudinally. During cell cycle progression sister *oriC*-ParB complexes segregate and move towards septal positions, whereby up to 4 ParB foci in MMI and up to 6 ParB foci per cell could be detected microscopically. Overlapping C-periods allow for short doubling times of down to 63 min with a new round of replication initiating 28 min after cell division and the ongoing replication event terminating 15 min later. Cells with long doubling times of 130 min possess a short time interval of seven min between cell division and replication initiation (B-period) and fully replicate both chromosome copies once per generation. In both, the fast and the slow growth model cells are diploid, since newly replicated sister chromosomes will only be separated by cell division in the daughter generation. Conclusively, cell cycle models suggested here describe overlapping replication cycles in combination with two sets of chromosomes in *C. glutamicum*.

**Figure 6.**
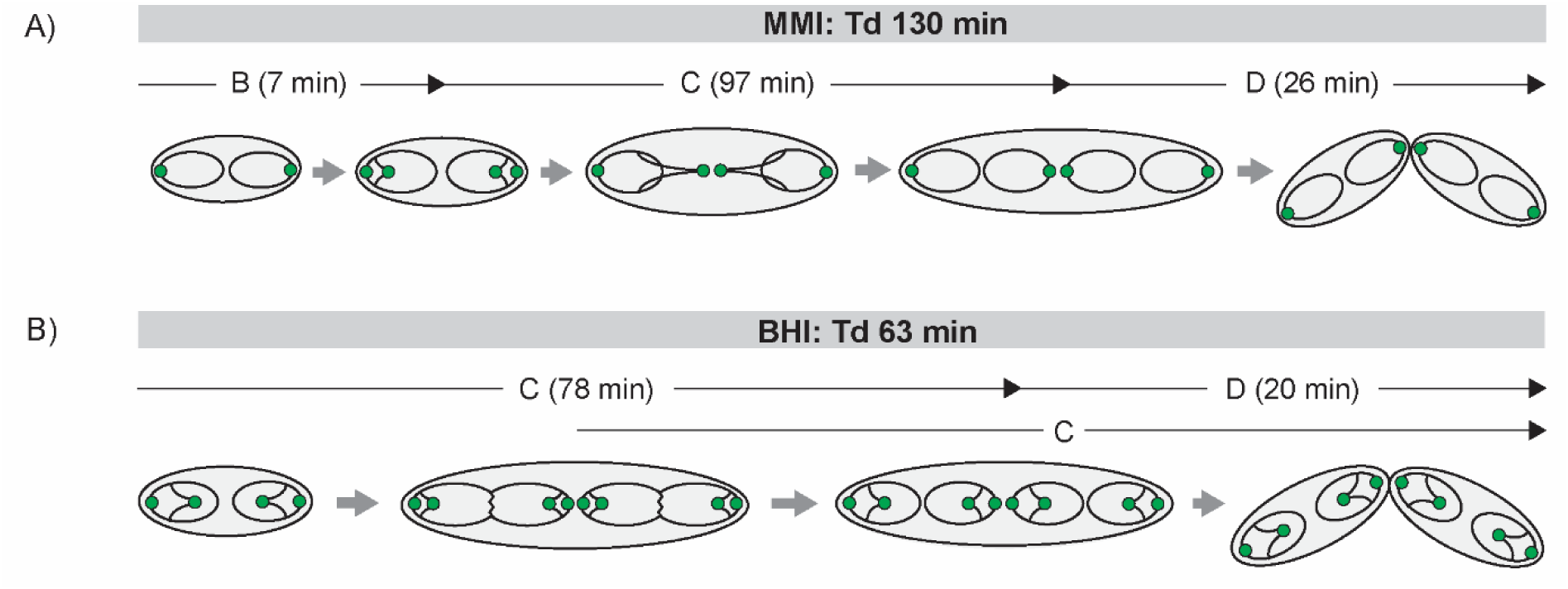
Spatiotemporal chromosome organization of *C. glutamicum*. Chromosomes are depicted as black lines with *oriCs* as green circles. In newborn cells two initial *oriC* locate close to the poles. Upon initiation of a new round of replication sister origins segregate non-synchronously from polar ParB-eYFP cluster and move towards midcell where a new septum is formed. Notably, stages with single chromosomes per cell are absent. **A)** Cell cycle of slow growing cells in MMI medium. A short B period is followed by C-and D-periods; replication takes place within one generation. **B)** Chromosome organization during fast growth in BHI medium. Multifork replication allows for short doubling times with a second round of replication starting after the first half of the cell cycle, around 15 min before the previous one terminates.

## Discussion

The bacterial cell cycle has been analyzed in few model organisms in the past. A hallmark of fast growing species such as *E. coli, B. subtilis* is the initiation of new rounds of DNA replication prior to replication termination and cytokinesis (15,16). This process has been termed multi-fork replication. In slow growing species, or species with asymmetric division such as *C. crescentus* and *M. smegmatis* C-periods are not overlapping (13,14). The increasing knowledge about bacterial cell biology has made clear that even fundamental cell processes such as cytokinesis and DNA replication might be organized and regulated in a far higher diversity than initially thought (76,77). We have therefore analyzed the cell cycle in *C. glutamicum* under various growth conditions and different growth rates. *C. glutamicum* emerges as a model organism for apical cell growth that is characteristic for actinomycetales (61).

Analysis of ParB foci in growing *C. glutamicum* cells revealed that under all culture conditions a ParB focus is stably attached to each pole, suggesting an *ori-ter-ter-ori* orientation of the chromosomes (Fig. 2 & 3). Interestingly, newly replicated origins segregate to midcell were they remain until cytokinesis is completed and they stay tethered to the newly forming cell poles (Fig. 3). Up to now it is unclear by which molecular mechanism the placement of ParB and the chromosomal origin is directed. Earlier work has shown that ParB and the cell division protein FtsZ interact (51). Interaction of ParB with the divisome might be a plausible explanation for the observed localization pattern. Deletion of ParA completely abolishes the directed ParB segregation, confirming earlier results from our group (51). Thus, unlike in other species with polar origin localization, such as *C. crescentus* and *V. cholerae*, the newly replicated origin is not migrating to the opposite pole (32,78). This mode of segregation in *C. glutamicum* is compatible with the observation that both cell poles are constantly occupied with a ParB-*oriC* complex, suggesting that even newborn cells contain at least two chromosomes and, hence, are diploid. Due to variable cohesion times of sister chromatids (Fig. 4C) the number of ParB foci does not necessarily reflect the number of origins and may lead to an underestimation of origins. The existence of several chromosomes is in line with the observation of multiple replication forks. During replication two or more replication forks can be visualized as judged by fluorescently labeled sliding clamp DnaN (Fig. 4). Localization of origins in *C. glutamicum* is therefore different from the situation in the closely related *M. smegmatis.* For *M. smegmatis* a replication factory model has been proposed in which a central localized origin is replicated and the newly replicated origins are segregated towards the cell pole while the replisome remains in the cell center (72). In contrast, we show by time lapse analysis that replication forks originate close to the cell pole and migrate towards midcell in *C. glutamicum* (Fig. 4E, Fig. S3). Thus, this organism does not have a static replication factory and, hence, is not consistent with the replication factory model proposed for *B. subtilis* (79). We also observed that both origins initiate replication around the same time. Further support for diploidy stems from flow cytometry data (Fig. 5). In replication run-out experiments cells contain a minimum of two and up to twelve chromosomes, being a clear indication of multiple initiation during fast growth conditions (growth rate above 0.6 h^-1^). Although presence of multiple chromosomes per cell has been suggested before (80), only single cell analysis unambiguously supports the diploidy of *C. glutamicum* cells. The simultaneous presence of two polar chromosomes is in stark contrast to findings recently reported for *Mycobacterium* species (72,73). Corynebacteria and Mycobacteria are closely related and, hence, it comes as a surprise that cellular organization of their chromosomes might be different. The constant diploidy of *C. glutamicum* could be a consequence of their environmental life style. Many soil bacteria have elaborated sophisticated methods to counteract various environmental stresses such as desiccation, nutrient shortage or exposure to DNA damaging agents. In fact, nucleic acids are prominent targets for desiccation induced damage (81). Cells carrying two or more chromosome equivalents will increase the change for correct DNA repair based on homologous recombination. In line with this hypothesis survival rates of coryneform bacteria are known to be high under various stresses including desiccation (82-84). Analyses of long-term preservations of microbial ecosystems in permafrost demonstrate that Corynebacteria are dominating older sediments (84).

In summary, we have provided detailed analyses of the cell cycle of *C. glutamicum* at different growth rates (Fig. 6). Data presented here point to a unique and so far undescribed cell cycle regulation with two polar attached chromosomes that simultaneously initiate replication. At fast growth conditions new rounds of replication can be initiated before the previous round is complete, similar to multi-fork replication. In contrast to other bacteria with polar oriented chromosomes, such as chromosome I from *V. cholerae* or *C. crescentus, C. glutamicum* cells contain two copies of the chromosomes and segregate the newly replicated origins only to midcell. The elucidation of the corynebacterial cell cycle is important to fully understand growth behavior and homologous recombination in this medically and industrially relevant genus.

## Acknowledgements

The authors are thankful to Dr. Andreas Brachmann (LMU) for help with the genome sequencing. We thank Dr. Stephan Gruber (Max-Planck-Institute for Biochemistry) for stimulating discussions and suggestions. This work was funded by grants from the Deutsche Forschungsgemeinschaft (BR2815/6-1) and the Ministry of Science and Education (BMBF: 031A302 e:Bio-Modul II: 0.6 plus).

